# Cell-type Annotation with Accurate Unseen Cell-type Identification Using Multiple References

**DOI:** 10.1101/2022.11.17.516980

**Authors:** Yi-Xuan Xiong, Meng-Guo Wang, Luonan Chen, Xiao-Fei Zhang

**Affiliations:** School of Mathematics and Statistics, Central China Normal University, Wuhan, China; Hubei Key Laboratory of Mathematical Sciences, Central China Normal University, Wuhan, China; State Key Laboratory of Cell Biology, Shanghai Institute of Biochemistry and Cell Biology, Center for Excellence in Molecular Cell Science, Chinese Academy of Sciences, Shanghai, China; School of Life Science and Technology, ShanghaiTech University, Shanghai, China; Key Laboratory of Systems Health Science of Zhejiang Province, Hangzhou Institute for Advanced Study, University of Chinese Academy of Sciences, Chinese Academy of Sciences, Hangzhou, China; Guangdong Institute of Intelligence Science and Technology, Hengqin, Zhuhai, Guangdong, China

## Abstract

The recent advances in single-cell RNA sequencing (scRNA-seq) techniques have stimulated efforts to identify and characterize the cellular composition of complex tissues. With the advent of various sequencing techniques, automated cell-type annotation using a well-annotated scRNA-seq reference becomes popular but relies on the diversity of cell types in the reference. There are generally unseen cell types in the query data of interest because most data atlases are obtained for different purposes and techniques. When annotating new query data, identifying unseen cell types is fundamental not only for improving annotation accuracy but also for novel biological discoveries. Here, we propose mtANN (multiple-reference-based scRNA-seq data annotation), a new method to automatically annotate query data while accurately identifying unseen cell types with the aid of multiple references. Key innovations of mtANN include the integration of deep learning and ensemble learning to improve prediction accuracy, and the introduction of a new metric defined from three complementary aspects to distinguish between unseen cell types and shared cell types. In addition, a data-driven method is provided to adaptively select threshold for unseen cell-type identification. We demonstrate the advantages of mtANN over state-of-the-art methods for unseen cell-type identification and cell-type annotation on two benchmark dataset collections, as well as its predictive power on a collection of COVID-19 datasets. The source code and tutorial are available at https://github.com/Zhangxf-ccnu/mtANN.

**Author summary:** Single-cell transcriptomics is rapidly advancing our understanding of the cellular composition of complex tissues and organisms. With the advent of various sequencing techniques, automatic cell-type annotation using well-annotated single-cell RNA sequencing (scRNA-seq) references has become popular. Compared with unsupervised cell-type annotation methods, it can be more easily applied to different data, saving labor and time costs. However, it relies on the diversity of cell types in the reference so there are generally unseen cell types in the query data. These unseen cell types need to be identified when annotating new sequencing data not only for improving annotation accuracy but also for novel biological discoveries. To address these issues, we propose mtANN, a new method to automatically annotate query data while accurately identify unseen cell types with the help of multiple references. We demonstrate the annotation performance of mtANN in PBMC and Pancreas collections when different proportions of unseen cell types are present in the query dataset. We also verify the practical application of mtANN in a collection of COVID-19 datasets for patients with different symptoms. When there are unseen cell types in the query dataset, mtANN is able to identify the unseen cell types and accurately annotate the shared cell types, especially the two cell types that are biologically similar.

## Introduction

Single-cell RNA sequencing (scRNA-seq) technologies can measure the gene expression profile of a sample at single-cell resolution. Their recent advances have stimulated efforts to identify and characterize the cellular composition of tissues, revolutionizing the understanding of the heterogeneity of complex tissues. With the various sequencing technologies, like 10x Genomics Chromium, Drop-seq, and Smart-seq2, having emerged, understanding the complex tissues has turned into a task of cell-type annotation for new sequencing data [1–3].

There are two typical solutions for cell-type annotation tasks. One of the solutions is to unsupervised cluster cells into groups based on the similarity of their gene expression profiles, and annotate cell populations by assigning labels to each cluster according to cluster-specific marker genes [4–8]. However, such methods require extensive literature review and manual testing of various combinations of marker genes, which is not only time-consuming but also not reproducible across different experiments within and across research groups [9, 10]. Another solution is to learn the intrinsic relationship between gene expression profiles and cell types based on a well-annotated reference atlas, and transfer the learned relationship to query data for cell-type annotation. There are two main types of approaches to this reference-based strategy, one is to learn the similarity between the reference atlas and the query data based on statistical metrics, as the basis for cell-type label transfer [11–14]. The other is to model a classifier on the reference atlas, which can make predictions directly on the query data [15–19]. The reference-based method can avoid manual selection of marker genes, and the trained classifier can be used for any new query data, providing convenience for practical applications.

Previous reference-based methods rarely take into account the following two issues. The first one is the selection of the reference atlas. Noise from the reference data and incorrectly annotated cell types may lead to inaccurate annotations on the query data, and the selection of input features of the classification model also has an impact on the annotation performance of different methods [20, 21]. This issue is expected to be partially addressed by integrating multiple well-annotated reference datasets and multiple gene selection methods [22–25], where an appropriate integration strategy is needed. Most of previous related methods first integrate multiple well-labeled datasets into a comprehensive reference atlas, and then annotate the cell types of the query data according to the integrated reference atlas, which makes the annotation results dependent on the removal of batch effects between different reference datasets [26, 27]. Another issue is that there is a difference in the joint distribution of gene expression and cell type between the reference and query datasets due to the difference in the marginal distributions. Distributional differences in gene expression, known as batch effects, have been extensively addressed in previous studies [28, 29], while differences in the distribution of cell types have been rarely considered. Discrepancies in cell types indicate that there are cell types in the query data that are not seen in the reference atlas, which can be called “unseen” cell types. Unseen cell types may suggest new biological discoveries that cannot be neglected. Additionally, ignoring the presence of unseen cell types biases the classifier learned on the reference atlas to known cell types, resulting in false predictions on the query data. These two issues are potentially related. Integrating multiple reference datasets can enrich the cell type information of reference data, but how to integrate reference datasets containing different cell types is difficult. In addition, how to identify cell types in the query data that are not seen in the reference data also requires effective methods.

In order to address the above two issues, we propose mtANN (multiple-reference-based scRNA-seq data annotation), a novel method that automatically identifies unseen cell types while accurately annotating query dataset by integrating multiple well-annotated scRNA-seq datasets as references. The main idea of mtANN is first to learn multiple deep classification models from multiple reference datasets to obtain multiple prediction results. These results are then used to vote on metaphase annotations and to compute metrics from three complementary aspects to identify unseen cell types. Final annotations are made based on metaphase annotation and unseen cell type identification results. mtANN has the following characteristics: (i) it utilizes the diversity of multiple reference datasets and avoids the selection of a single reference dataset; (ii) it combines the ideas of deep learning and ensemble learning to improve prediction accuracy; (iii) it proposes a new metric from three complementary aspects to measure whether a cell belongs to an unseen cell-type; and (iv) it introduces a new data-driven approach to automatically determine thresholds for unseen cell type identification. We benchmark mtANN using two collections of benchmark datasets, each from different tissues, sequencing technologies, and containing different cell types. We prepare a total of 75 benchmark tests, including annotations across different technologies and unseen type belonging to different cell types. We also use a COVID-19 dataset and prepare a total of 249 tests to assess the performance. Experimental results demonstrate that mtANN outperforms state-of-the-art methods in both unseen cell-type identification and cell-type annotation.

## Results

### Overview of mtANN

The workflow of mtANN is illustrated in Fig 1 and S1 Text. To simultaneously annotate the query data and identify unseen cell types, mtANN first adopts eight gene selection methods to generate a series of subsets that retain distinct genes for each reference dataset (Fig 1A). This step facilitates the detection of biologically important genes and increases data diversity for effective ensemble learning. Second, mtANN trains a series of neural network-based deep classification models based on all reference subsets. These base classification models characterize different relationships between gene expression and cell types which are complementary in identifying unseen cell types. (Fig 1A). Third, mtANN annotates the query dataset by integrating the outputs of all base classification models (Fig 1B). Fourth, mtANN defines a new metric from three complementary aspects to identify cells that may belong to unseen cell types (Fig 1C). Finally, we provide a data-driven method for threshold selection, which helps to mark cells belonging as “unassigned” (Fig 1C).

**Fig 1.**
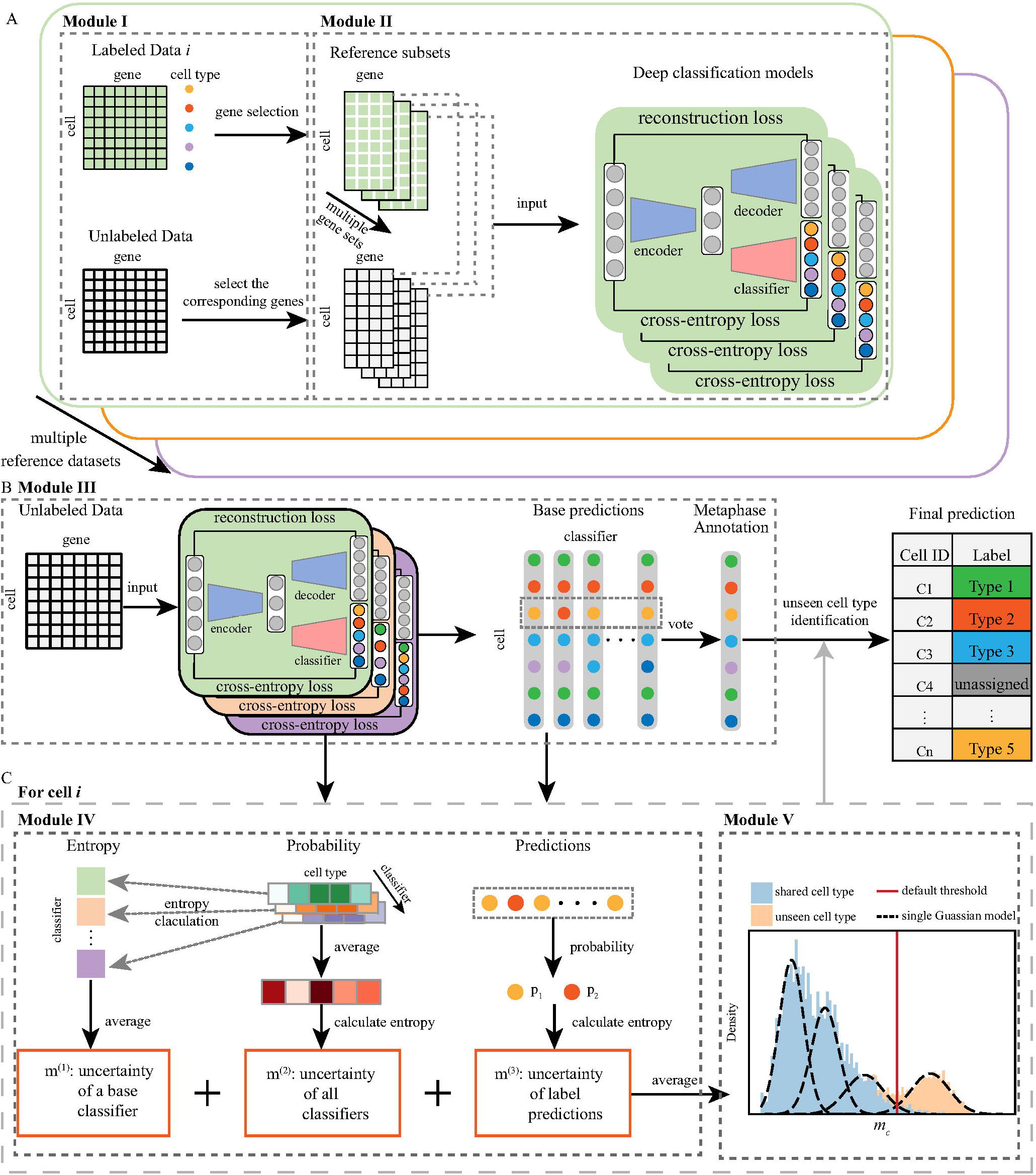
Overview of mtANN. (A) The training process of mtANN includes two modules: gene selection (Module I) and deep classification model training (Module II). The labeled data *i* is used as an example. In Module I, eight gene selection methods are applied on data i, obtaining multiple reference subsets. The gene sets selected by the eight gene selection methods intersect with all the genes in the query dataset, determining the input genes of multiple deep classification models. In Module II, pairs of reference subset and query dataset after gene selection are used as input to train each deep classification model. We conduct theses two modules for every labeled data, thus obtaining multiple deep classification models. (B) The prediction process of mtANN (Module III) first makes predictions for the query data based on deep classification models learned by Module II and then conducts a majority vote to obtain a metaphase annotation. (C) Unseen cell type identification process, consisting of two modules: quantifying the likelihood of a cell belonging to an unseen cell type (Module IV) and using a data-driven threshold determination method to identify unseen cell types (Module V). In Module IV, we define an unseen cell type identification metric by averaging three uncertainty measures calculated from the results obtained from III. In Module V, we derive a new a data-driven method based on Gaussian mixture model to determine the threshold for unseen type identification. If a cell is identified as belonging to an unseen cell type, mtANN annotates it as “unassigned”; otherwise mtANN annotates it as the result of module III.

### Validation of the rationale of mtANN

mtANN integrates multiple well-annotated scRNA-seq datasets as references and applies eight gene selection methods to select informative genes. To validate the effectiveness of integrating multiple reference datasets and gene selection methods, we use two collections of datasets from two tissues: peripheral blood mononuclear cells (PBMC) collection which contains 7 datasets sequenced by 7 different technologies [20] and Pancreas collection containing 4 datasets sequenced by 4 different technologies [30–33] (Methods Datasets section). In each collection, we alternately select one dataset as a query dataset and the rest as reference datasets. The eight gene selection methods denoted as DE, DV, DD, DP, BI, GC, Disp, and Vst (Methods Gene selection section) are applied to these reference datasets separately, obtaining multiple reference subsets. Classification models trained on a single reference subset are compared with mtANN that integrates the results from different models to show the effectiveness of ensemble learning.

As an illustrative example, we first demonstrate the effectiveness of mtANN using the “Celseq” dataset from the PBMC collection and the “Baron” dataset from the Pancreas collection. When “Celseq” is used as the query data, the remaining “Drops”, “inDrop”, “Seq-Well”, “Smart-seq2”, “10X v2”, and “10X v3” are all used as reference datasets. In the Pancreas collection, the “Muraro”, “Segerstolpe”, and “Xin” are used as reference datasets while “Baron” is used as the query dataset.To show the difference in gene selection methods, we present the performance of different gene selection methods with different colored points. In Fig 2, it is clear that the red line is higher than all the points, indicating that mtANN’s strategy of integrating all reference datasets and gene selection methods is consistently superior to a single classification model. It is worth noting that the performance of different gene selection methods varies across reference datasets, and no single gene selection method always wins on all datasets. Similar results are also observed in experiments with other datasets from both data collections (S1 Fig). These results demonstrate the necessity and effectiveness of integrating multiple reference datasets and gene selection methods to annotate cell types.

**Fig 2.**
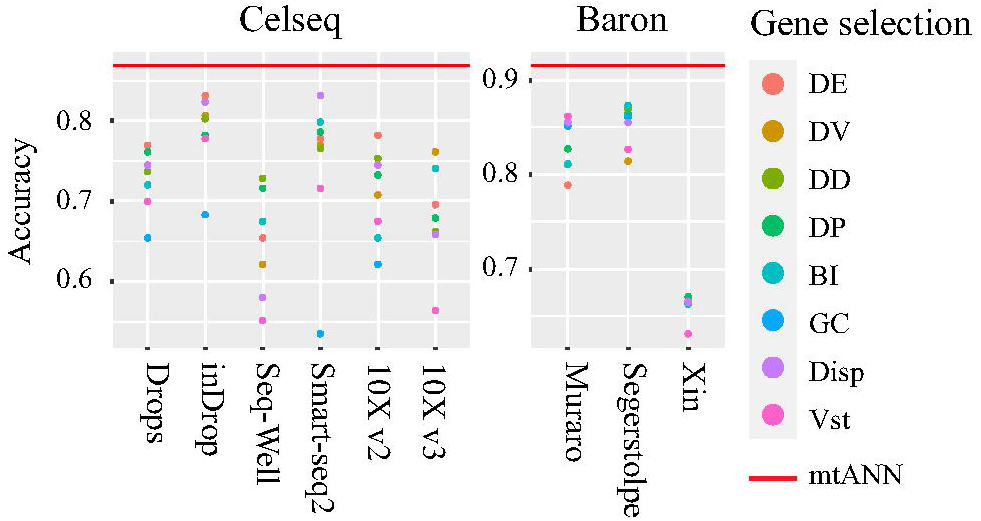
Accuracy comparison between mtANN and each base classification model. The “Celseq” dataset in the PBMC collection and the “Baron” dataset in the Pancreas collection used as query datasets are displayed. In each plot, each column represents a reference dataset. Each point is the performance of a base classification model, and points of different colors represent different gene selection methods. The red line indicates the performance of mtANN that integrates different reference datasets and gene selection methods.

### Benchmarking mtANN for unseen cell-type identification

mtANN is specifically designed for unseen cell type identification when annotating cell types. To demonstrate its ability in identifying unseen cell types, we also use the two data collections: PBMC and Pancreas. Within each data collection, each dataset is alternately used as a query dataset and the rest are used as reference datasets. To simulate unseen cell type in the query dataset, we perform a leave-one-cell-type-out setting in every references-query pair. In doing so, we obtain a total of 50 tests in the PBMC collection and 25 tests in the Pancreas collection (for details, please refer to S2 Fig and S5 Table-S6 Table). We compare mtANN with several existing popular methods, including scmap-clust, scmap-cell [11], Seurat v3 [12], ItClust [15], scGCN (entropy), scGCN (enrichment) [16], and scANVI [17] (S1 Text Methods for benchmark section) as they also provide metrics for unseen cell-type identification. To evaluate each method in distinguishing unseen cell types from shared cell types, we compare the performance of each method in terms of AUPRC scores (S1 Text Performance assessment section).

The results are presented in Fig 3A and S3 Fig. Fig 3A shows that mtANN is superior to the compared methods when the “10X v3” dataset is the query. The results on other datasets also show the superior performance of mtANN (S3 Fig). Across all the experiments, we count the number of times each method ranks first in terms of AUPRC scores. We observe that the performance of scmap-clust, ItClust, and scGCN (enrichment) vary widely between different data collections (S4 Fig). They may rank first in some datasets, but have a large performance drop in others. This may be due to the difference in the distribution of cell types in the query and reference datasets. Take ItClust as an example, missing cell types in the reference data may lead to misalignment of cell labels and clusters, resulting in over-fitting of the model. mtANN effectively addresses this issue by borrowing complementary information between different reference datasets to define the uncertainty of cell annotation at different aspects, thereby accurately distinguishing between shared cell types and unseen cell types (Fig 3B and S5 Fig-S7 Fig).

**Fig 3.**
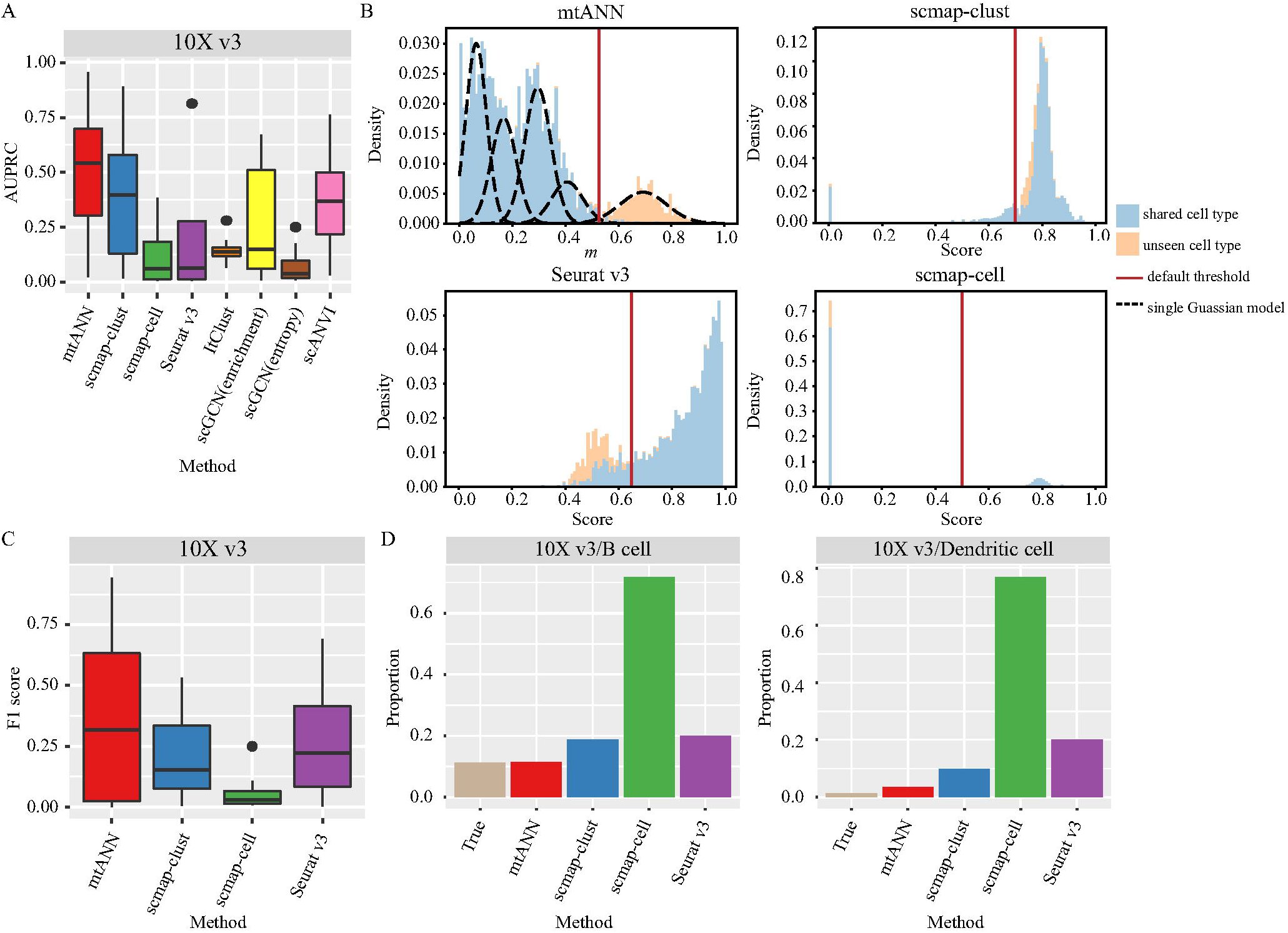
The performance in unseen cell-type identification. (A) The boxplot of the AUPRC for mtANN and other methods when “10X v3” in PBMC collection is the query dataset. (B) The distribution of each metric that measures cell prediction uncertainty when the “10X v3” dataset in the PBMC collection is the query dataset and “B cell” is the real unseen cell type. The color of the histogram distinguishes unseen cell types from shared cell types. The black dotted line represents the subpopulations of the Gaussian mixture model fitted by mtANN. The red solid line represents the default threshold selected by each method. Cells with a metric greater than the threshold are identified as “unassigned” in mtANN and cells with a score less than the threshold are identified as “unassigned” in scmap-clust, scmap-cell, and Seurat v3. (C) The boxplot of the F1 score for mtANN and other methods when “10X v3” in PBMC collection is the query dataset. We use the default threshold provided by each method to select “unassigned” cells. (D) The bar plot of the proportion of unseen cell types and “unassigned” cells predicted by each method when the “10X v3” dataset in the PBMC collection is the query dataset. The name of each plot indicates the real unseen cell type.

Another important issue in the identification of unseen cell types is the choice of thresholds. Most methods for identifying unseen cell types choose a fixed threshold (scmap) or a fixed ratio (Seurat v3) as the threshold, which may not generalize well on new datasets. To test the performance of the threshold selection methods, mtANN is compared with scmap-clust, scmap-cell and Seurat v3, which have provided threshold selection methods, in terms of F1 score. The results are presented in Fig 3C and S8 Fig. It can be seen that mtANN performs better than other methods in most cases, scmap-cell performs the worst in PBMC collection, and the relative performance of scmap-clust, scmap-cell and Seurat v3 varies by dataset. To further investigate the reasons for the differences in the performance of these methods, we compare the proportion of true unseen cell types with the proportion of cells identified as “unassigned” by each method (S9 Fig). We take an experiment where “10X v3” is the query dataset, as an example (Fig 3D). When the unseen cell type is B cell, the true proportion of the unseen cell type is 11%, and the proportion of cells predicted by mtANN as “unassigned” is close to 11%. While the “unassigned” cell proportion predicted by scmap-cell is much higher than the true proportion, and the proportion of “unassigned” cells identified by Seurat v3, fixed at 20%, is also higher than the true proportion. When the unseen cell type is Dendritic cell, the proportion of unseen cell types is small. The proportion of cells predicted by mtANN as “unassigned” decreases and is close to the true proportion, while the proportions of cells predicted by scmap-clust, scmap-cell, and Seurat v3 as “unassigned” are much higher than the true proportion. With default thresholds, we count the number of times each method ranks first in terms of F1 score across all the experiments, and find that mtANN is consistently able to accurately identify unseen cell types when the proportion of unseen cell types is varied (S4 Fig).

### Benchmarking mtANN for cell-type annotation

In addition to identifying unseen cell types, annotation of new query data requires label annotation of cells belonging to shared types. To evaluate the performance of mtANN for annotating query datasets when there are unseen cell types, we also use the PBMC and Pancreas collections to conduct the experiments. For each experiment, one dataset is selected as the query dataset and the rest are used as reference datasets. Since the presence of unseen cell types may affect the annotation of shared cell types, we still use the leave-one-cell-type-out setting in each experiment. As threshold selection will affect the annotation accuracy of the query dataset, we conduct two ways to determine the thresholds: the real proportion of unseen cells and the default threshold provided by each method.

When using the real proportion of unseen cell types in the query dataset as the threshold to assign unseen cells, the annotation accuracy of mtANN and other methods are presented in Fig 4A and S10 Fig. It can be observed that in different experiments with different proportions of unseen cell types, mtANN always achieves higher annotation accuracy than other methods (S4 Fig).The performance of scmap, Seurat v3, and ItClust varies greatly across different experiments. This may be due to the presence of unseen cell types in the query dataset, leading to annotation bias towards shared cell types. To validate this, we calculate the Pearson correlation coefficient between the true proportional distribution of cell type in each experiment and the proportional distribution of the predicted results of each method (S11 Fig). The results show that mtANN has the highest correlation in both PBMC and Pancreas collections. We also use 10X v3 datasets being the query dataset and B cell being the real unseen cell type as an example. Fig 4B shows that the proportion of cell types obtained from mtANN’s prediction is more similar to the true proportion. In detail, we find that mtANN identifies most B cells as “unassigned”, whereas all other comparison methods annotate most B cells as a similar cell type (CD4+ T cells) as they are all derived from lymphoid progenitors (Fig 4C). For shared cell types, mtANN performs better at distinguishing the two monocyte subtypes, while scmap-clust, scmap-cell, and Seurat v3 tend to confuse CD16+ monocyte cells with CD14+ monocyte cells. scGCN (enrichment), scGCN (entropy), and scANVI fail to annotate monocytes and other rare cell types (Dendritic cell, Megakaryocyte, Natural killer cell, and Plasmacytoid dendritic cell). Overall, mtANN performs better than other methods in annotating new query data containing unseen cell types.

**Fig 4.**
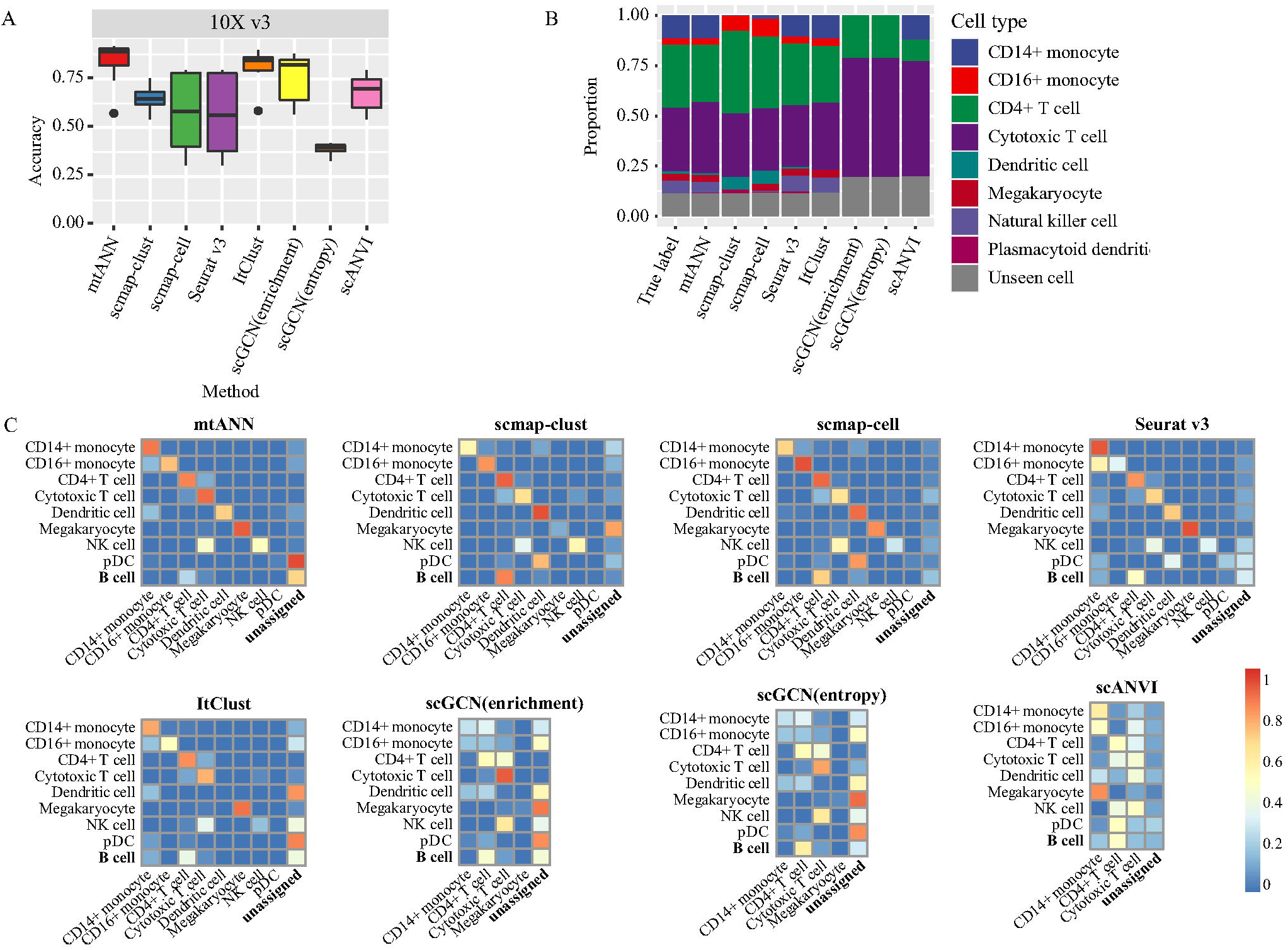
The performance in cell-type annotation when there are unseen cell types in query dataset. (A) The boxplot of the accuracy for mtANN and other methods when the “10X v3” dataset in the PBMC collection is the query dataset. (B) The bar plot of the real proportion of cell types and the proportion of cell types annotated by each method when the “10X v3” dataset in the PBMC collection is the query dataset and “B cell” is the real unseen cell type. (C) Heatmaps of the confusion matrices of mtANN and other methods when the “10X v3” dataset is the query dataset and “B cell” is the real unseen cell type. In a confusion matrix, the row and column names correspond to the true cell labels and the predicted cell labels of the query dataset, where the element represents the proportion of cells belonging to one cell type that is predicted to be of other cell types. NK cell and pDC are abbreviations for the Natural killer cell and the Plasmacytoid dendritic cell.

In reality, we cannot obtain the real proportion of unseen cell types, so the default threshold provided by each method is more practical and essential. When using the default method to select threshold, the prediction accuracies of mtANN, scmap-clust, scmap-cell and Seurat v3 in all the tests are presented in S12 Fig. We can observe that the accuracy of mtANN is higher than those of the compared methods when “Celseq”, “Drops”, “inDrop”, “Smart-seq2”, “10X v2”, and “10X v3” are evaluated as the query datasets (S12 FigA). S12 FigB shows that mtANN also has the best performance when “Baron human” and “Xin” are used as the query datasets. In addition, the result of mtANN at the default threshold is similar to the result at the actual proportion (S13 Fig), indicating that the threshold selected by mtANN is comparable to the actual proportion of unseen cells.

### Cell-type annotation of COVID-19 patients with different symptoms

Coronavirus disease 2019 (COVID-19) has caused more than 647 million infections and more than 6.6 million deaths, according to World Health Organization (WHO) statistics as of December 16, 2022. It is thus important to annotate the cell types of the sequencing data from COVID-19 patients for understanding the disease mechanism. With many scRNA-seq data from COVID-19 patients available, we select the study of COVID-19 that offers a comprehensive immune landscape [34], including 284 samples from 196 COVID-19 patients and controls to assess the performance of mtANN on real data. We use the dataset from PBMC cells in the COVID-19 dataset as the query datasets and the PBMC collection we used above [20] as references to evaluate the performance of mtANN and other methods.

We group the cells according to samples’ id, resulting in 249 query datasets. mtANN is compared with scmap-clust, scmap-cell, and Seurat v3 under the default threshold parameters for identifying unseen cell types. The accuracies of mtANN and other methods on the 249 query datasets are presented in Fig 5A. It can be seen that the accuracies of mtANN for patients with different symptoms are higher than other methods, and scmap-cell suffers a decrease. We further conduct a one-to-one comparison and find that mtANN significantly (two-sided paired Wilcoxon test, p-value < 0.01) outperforms the compared methods (Fig 5B). We compare the composition of cell types between patients with different symptoms and find that the proportion of B cells increases in patients with severe symptoms, and the percentage of dendritic cells and T cells decreases, particularly in patients with severe symptoms (Fig 5C), which is consistent with the lymphopenia phenomenon previously reported [35]. We also find that the percentage of megakaryocyte and CD14+ monocyte elevates in patients with severe symptoms, which is agreement with the original study [34]. Compared with scmap-clust and Seurat v3, mtANN can more accurately reflect the difference in the proportion of Dendritic cell and Megakaryocyte cells between different populations, which is instructive for the study of the development process of COVID-19.

**Fig 5.**
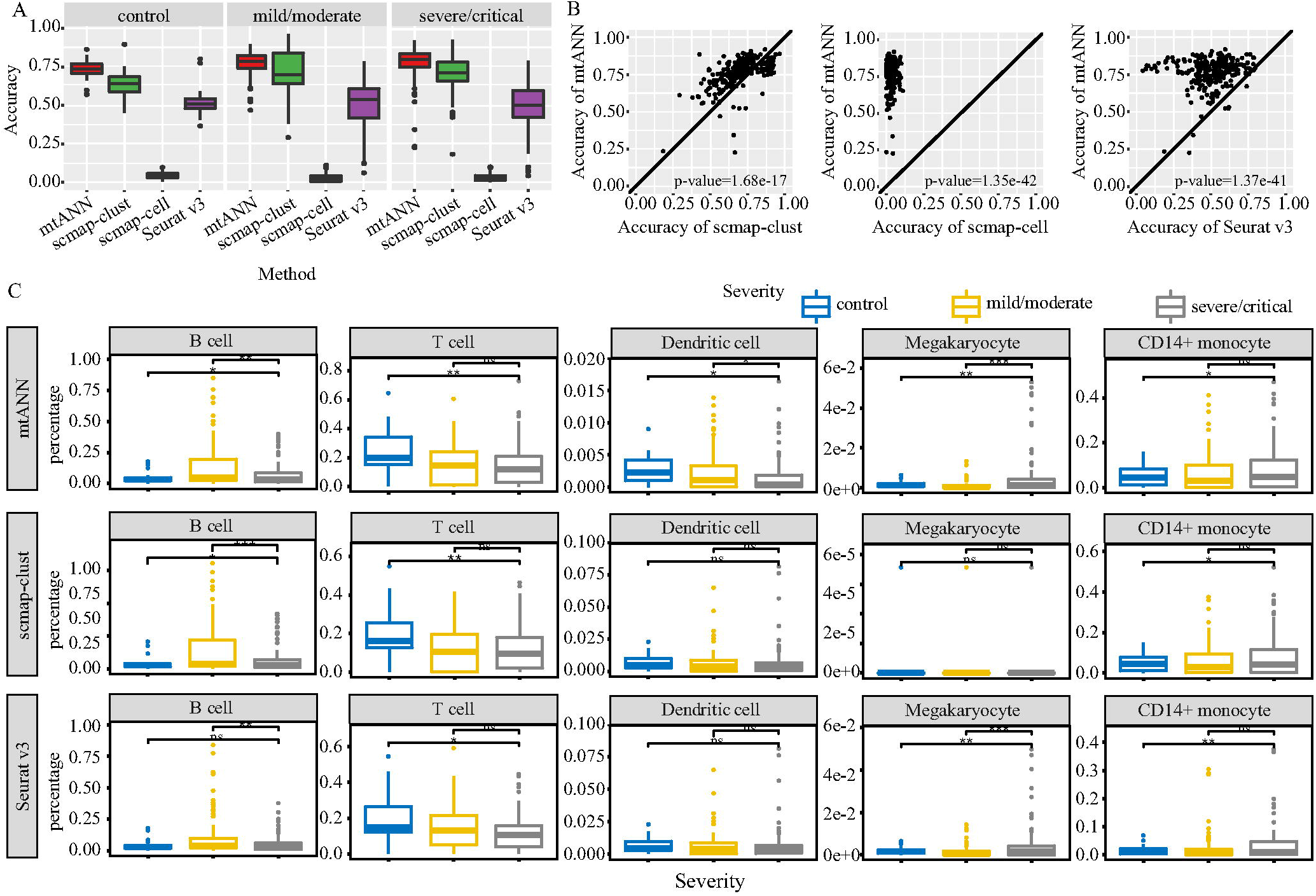
The application of different methods on COVID-19 dataset. (A) The boxplots of the accuracy of different methods on samples in different symptoms. (B) One-to-one comparison between mtANN and scmap-clust, scmap-cell, and Seurat v3. Each point represents a query dataset. P-values of two-sided paired Wilcoxon signed-rank tests used to test for significance of differences in performance are reported. (C) The boxplots of the compositions of B cells, T cells, Dendritic cells, Megakaryocyte cells, and CD14+ monocyte between samples with different symptoms. The significance of the two-sided T-test is represented by stars where one star, two stars and, three stars mean the corresponding p-value less than 0.05, 0.01 and, 0.001, respectively, and ns means the corresponding p-value greater than 0.05.

## Discussion

With the development of single-cell sequencing technology, traditional unsupervised clustering-based cell-type annotation methods are difficult to adapt to rapidly generated datasets since they are time-consuming [36, 37]. Another method for automatic cell-type annotation based on a reference atlas has been widely studied, but these methods are rarely able to discover unseen cell types in the query data [23, 38, 39]. The identification of unseen cell types may lead to new biological discoveries, while erroneous identification may lead to missing new biological discoveries or lead improper biological conclusions. Only some of the previous methods for automatically annotating cell types address the problem of identifying unseen cell types, and all of them only set a default threshold instead of proposing a methodology to automatically select a threshold. The choice of threshold can determine the accuracy and usability of the method.

In this study, we propose a novel ensemble learning-based cell-type annotation method, mtANN, to annotate cell-type labels for a query dataset automatically. Our method mainly has three innovations: (i) it integrates multiple reference datasets not only to enrich cell types in the reference atlas but also to provide complementary information to annotate cell types; (ii) it proposes a new metric from three complementary aspects to effectively measure whether a cell belongs to unseen cell types; and (iii) it proposes a data-driven approach to adaptively determine the threshold for unseen cell-type identification. Through the 75 sets of experiments we constructed, we demonstrate the annotation ability of mtANN for new sequencing data, and validate that mtANN can also accurately distinguish between unseen cell types and shared cell types when the proportion of unseen cell types in the query dataset varies. In particular, mtANN also has excellent discrimination between two similar cell types in the shared cell types. The application on the real data verifies the annotation performance of mtANN for COVID-19 patients, and its annotation results can show the difference in the proportions of different immune cells between different populations. Our comprehensive benchmark and application on an extensive set of publicly available benchmark datasets indicate that mtANN has achieved state-of-the-art performance for unseen cell-type identification and cell-type annotation in the meantime.

There may be two limitations in integrating multiple reference datasets for unseen cell-type identification that we have not addressed well in this work. One limitation of our method is the inconsistent terminology of cell types across different reference datasets. In this work, we avoid this problem by manually checking the cell-type annotations. For example, the cell type “PP” in the Xin dataset is changed to “gamma”, as “gamma” is the name used by all other datasets. Several approaches can be attempted in the future to match cell types between datasets, such as matching based on marker genes of cell types or matching by mutual prediction between datasets. In this work, we only mark the cells that are considered to belong to unseen cell types as “unassigned”. Thus, another limitation is that we do not provide a further biological interpretation of these cells. A straightforward way to determine the identities of these cells is to use unsupervised annotation methods. In addition, integrating Cell Ontology [40, 41] into the method may be instructive for annotation of “unassigned” cells. For example, when Plasmacytoid dendritic cells are absent from reference dataset, they can be assigned to supertypes of Dendritic cells with the help of Cell Ontology. In the future, we will extend our method to implement this functionality.

## Materials and methods

### Module I: Gene selection

mtANN uses five supervised gene selection methods collected by scClassify [23] and three widely used unsupervised methods for highly variable gene selection (for details, please refer to S1 Text). For each reference dataset (*X^r_i_^, Y^r_i_^*), we adopt the eight gene selection methods to pick genes from different perspectives, including Limma, Bartlett’s test, Kolmogorov-Smirnov test, Chi-squared test, Bimodality index, Gini index-based clustering [42], Dispersion and Variance-stabilizing transformation [12]. The first five methods select differentially expressed genes (DE), differential variable genes (DV), differentially distributed genes (DD), differentially proportioned genes (DP), and bimodally distributed genes (BI) respectively, and the latter three methods pick highly variable genes based on Gini index-based clustering (GC), dispersion (Disp), and variance (Vst). We index these gene selection methods using *j* = 1,…, 8. Let *G^r_ij_^* denotes the gene set selected by the *j*-th gene selection method for the *i*-th reference dataset, where *r_ij_*, *i* = 1,…, *M, j* = 1,…, 8 is the index of reference subsets, and *G^q^* denote all genes in the query dataset. In doing so, we can obtain 8*M* reference subsets to expand the diversity of references. In each reference subset, we can train a deep classification model with *G^r_ij_^* ⋂ *G^q^* as the input features. We denote *X^r_ij_^* and *X^q_ij_^* as the gene expression matrix after gene selection for the *ij*-th reference subset and query dataset. For convenience, we still denote the preprocessed data by *X^r_ij_^* and *X^q_ij_^*. Based on preprocessed data, we construct a dataset pair (*X^r_ij_^, Y^r_i_^, X^q_ij_^*), as the training dataset for the next step to train a base classification model.

### Module II: Deep classification model training

Based on each dataset pair (*X^r_ij_^, Y^r_i_^, X^q_ij_^*), we train a classification model based on deep learning. The classification model involves two components: the embedding component for extracting cell type-related features and the linear classifier layer for classification. Let *E^ij^* and *C^ij^* denote the embedding component and the linear classifier layer separately. The forward propagation result of the classification model after softmax transformation can be defined as 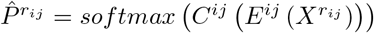, where 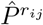 is an assignment probability matrix with rows representing cells and columns representing cell types. The (*c, k*)-th element of 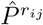 can be regarded as the predicted probability of cell *c* in *i*-th reference subset belonging to cell type *k*. The cross-entropy loss

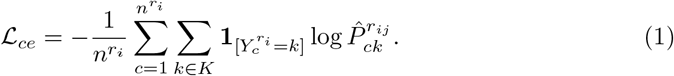

is used to train the classification model, where **1**_[.]_ denote the indicative function, and *n^r_i_^* is the number of cells in *i*-th reference dataset.

To enable the embedding component *E^ij^* to fully capture the characteristics of cells and make the classification model better fit the query dataset, we employ the embedding component as an encoder and use a mirror image of the embedding component as a decoder to construct an autoencoder. The reconstruction loss of cells both from the reference subset and the query subset is taken into consideration when training the classification model. Let *D^ij^* denote the decoder component. The forward propagation results of the autoencoder can be defined as 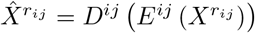 and 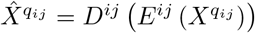, where 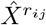 and 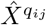 denote the reconstruction of *X^r_ij_^* and *X^q_ij_^* separately. The reconstruction loss is measured by the mean squared error, which can be formulated as

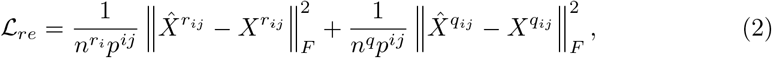

where *p^ij^* represents the number of genes in this dataset pair, and ||·||_*F*_ denote the Frobenius norm of a matrix.

Therefore, the final optimization problem for training the classification model for dataset pair (*X^r_ij_^, Y^r_i_^, X^q_ij_^*) can be written as

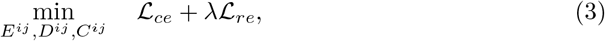

where λ is the tuning parameter and the default value is 1. Details of the neural network architecture, hyperparameter settings, and initialization can be found in S1 Text. For all the reference-query pairs, we can have 8*M* base classification models denoted by {(*E^ij^,C^ij^*)}_*i*=1,…,*M*,*j*=1,…,8_.

### Module III: Query dataset annotation

Based on one base classification model (*E^ij^, C^ij^*), we take the corresponding query subset *X^q_ij_^* as input. The forward propagation result along the model after softmax transformation can be formulated as 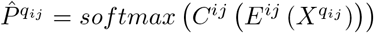. The (*c, k*)-th element of 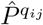 can be regarded as the predicted probability of cell *c* in the query dataset belonging to cell type *k*. For each cell in the query dataset, we obtain *q_ij_*-th base prediction label *Ŷ^q_ij_^* according to 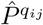. For cell *c*,

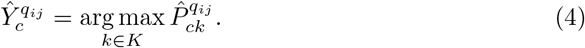

Then, based on the majority voting principle we integrate all these predictions for consensus annotation, denoted by *Ŷ^q^*. For cell *c*, we calculate

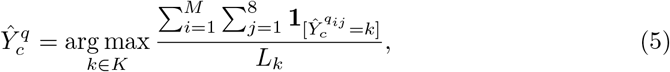

where 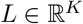, and *L_k_* indicates the number of reference subsets which contain cells belong to cell type *k*. The numerator represents the number of times that cell *c* is predicted to belong to cell type *k* across all base predictions and the denominator represents the number of reference subsets containing cell type *k*. The role of the denominator is to handle the situation where a cell is predicted as a single-reference-specific cell type and indeed belongs to that cell type in the query dataset. It is worth stating that the setting of the denominator increases the prediction probability of the single-reference-specific cell type, making full use of the diversity of the reference datasets. Details of the integration can be found in S1 Text.

### Module IV: Metrics for unseen cell identification

Since there is no training data in reference datasets for the unseen cell types, the predictions for the cells belonging to these cell types can be more uncertain. We define the uncertainty from two perspectives based on the outputs of all the classification models, including the intra-model perspective and inter-model perspective. For the former, no one cell type dominates the probability when making predictions based on a single classification model. For the latter, there is a large inconsistency among the predictions obtained by different classification models. Therefore, we design three entropy-based measures, denoted by *m*^(1)^, *m*^(2)^ and *m*^(3)^, to quantitatively characterize the uncertainty, where *m*^(1)^ is from the intra-model perspective, and *m*^(2)^ and *m*^(3)^ are from the inter-model perspective.

#### Intra-model measurement from each single classification model

The first metric *m*^(1)^ calculates the entropy of the probability that a cell belongs to different cell types by each classification model, and then averages these entropy values as a final uncertainty measure. Higher *m*^(1)^ indicates higher uncertainty. For cell *c*, this metric is defined as

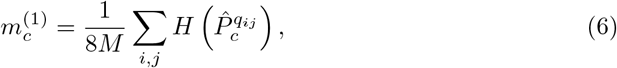

where *H*(·) represents the function to compute an entropy and is defined as 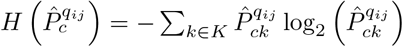. The larger 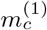 is, the more uncertain the predictions is, and thus the more likely the cell *c* is of unseen cell types.

#### Inter-model measurement from the overall output of classification models

The second measure *m*^(2)^ characterizes uncertainty from the inter-model perspective by first averaging the prediction probabilities of all classification models and then calculating the entropy. We compute the average of prediction probabilities *Q*^(2)^ as

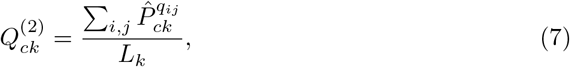

where 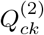 represents the average of the prediction probability that cell *c* belongs to cell type *k* across all classification models. Then, *Q*^(2)^ is transformed into a probability matrix 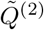 by dividing each value by the row sum. If the different base prediction labels for cell *c* are inconsistent, no one cell type dominates the probability in 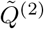. Therefore, we define *m*^(2)^ as the entropy of the average of the prediction probability to characterize inconsistency. For cell *c*, it is defined as

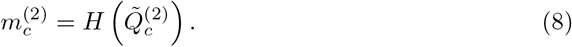

The larger 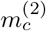 indicates less consistent predictions, and thus cell *c* is more likely to be of an unseen cell type.

#### Inter-model measurement from the overall hard-assignment labels

The third measure *m*^(3)^ is similar to *m*^(2)^. The difference is that it integrates the hard-assignment labels of all classification models rather than the prediction probabilities. Let *Q*^(3)^ denotes the integration result for this measure. The (*c, k*)-th element of *Q*^(3)^ is defined as

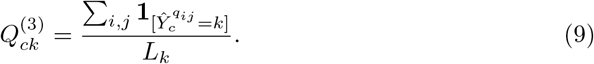

Then, as before, we transform *Q*^(3)^ into a probability matrix 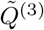 by dividing each value by the row sum. If the hard-assignment labels for cell *c* are inconsistent, then none of cell types dominate the row *c* of 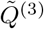. Similarly, we calculate the entropy to define *m*^(3)^, i.e., for cell *c*,

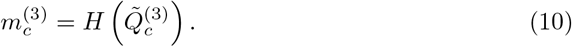

After obtaining the three complementary metrics, *m*^(1)^, *m*^(2)^ and *m*^(3)^, the values are scaled to [0, 1] linearly through Min-Max scaling separately, denoted by 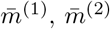 and 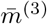. The ensemble uncertainty measure m is defined as the average of these three measures for each cell, i.e., for cell *c*,

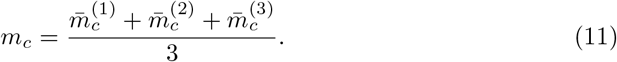

Generally, a larger value of *m_c_* indicates a higher probability that cell *c* belongs to an unseen cell type. Details of the calculation of each measurement can be found in S1 Text.

### Module V: Data-driven method for default threshold selection

Determining the threshold to distinguish cells belonging to unseen cell types remains subjective in previous studies, and a method to automatically determine the exact threshold is required. Instead of using a fixed value as a threshold as in previous studies, we provide a new method to automatically identify cells with higher uncertainty. We first fit a Gaussian mixture model to the uncertainty metric *m* with the number of mixture components *S* set to 1, 2,…, 5 and determined according to the Akaike information criterion (AIC). If the suitable value of S determined by AIC is 1, we consider that no cells are assigned as “unassigned”. Otherwise, all the cells are divided into different groups according to the posterior probability of the estimated Gaussian mixture model, and then the mean of the metric m of cells within each group is calculated. If there are groups with a mean greater than or equal to 0.6, these groups are considered to be uncertain groups. Meanwhile, the group with the largest mean is considered to be the uncertain group. All the cells in the uncertain groups are annotated as “unassigned”.

### Datasets

We use two collections of publicly available scRNA-seq datasets and a study of COVID-19 patients (S3 Table-S4 Table) varying from tissues (peripheral blood mononuclear cells (PBMC) and Pancreas), cell populations and sequencing technologies to benchmark mtANN and other methods.

The PBMC collection, including seven datasets curated from Butler et al. [43], are sequenced by Cel-seq, Drops, inDrop, Seq-Well, Smart-seq2, 10X v2, and 10X v3. The datasets are downloaded from https://doi.org/10.5281/zenodo.3357167 [20]. The Pancreas collection, including four datasets curated from Baron et al. [30], Muraro et al. [31], Segerstolpe et al. [32], and Xin et al. [33], are sequenced by inDrop, Cel-seq2, Smart-seq2, and SMARTer. We obtain all the datasets from https://hemberg-lab.github.io/scRNA.seq.datasets/human/Pancreas/. Following the study of scClassify [23], we manually check the cell-type labels that are provided by the original authors of each dataset and remove the cell types that are labeled as “unclear” in the Muraro dataset, “co-expression”, “not applicable”, “unclassified” and “unclassified endocrine” in Segerstolpe dataset, and “alpha.contaminated”, “beta.contaminated”, “delta.contaminated” and “gamma.contaminated” in Xin dataset.

The study of COVID-19 [34] provides a scRNA-seq atlas including 284 samples from PBMC, bronchoalveolar lavage fluid (BALF), sputum, and pleural fluid mononuclear cells (PFMCs) which is available at GEO database: GSE158055. In this study, we only take 249 of these samples from PBMC. We manually renamed CD8+ T cells to Cytotoxic T cells to be consistent with the previous PBMC collection.

For the PBMC and Pancreas data collections, we first remove cell types with less than 10 cells, then genes expressed in less than 100 cells are removed, and cells expressing less than 100 genes are later removed. These datasets are selected to be either the reference or the query datasets in the following experiments. For details about the reference and query datasets used in the benchmark tests, please refer to S5 Table-S6 Table.

### Data preprocessing

For each scRNA-seq dataset, preprocessing consists of four steps. Firstly, the library size normalization is performed, i.e., dividing the expression of each gene in a cell by the total expression of the cell and then multiplying it by a scale factor of 10000 in order to make the total expression values of all cells after being transformed the same. Secondly, logarithmic transformation is applied to the dataset to make each expression value *x* be log_2_ (*x* + 1). Thirdly, z-score standardization is performed for each gene so that the mean of each gene on all cells is equal to 0 and the standard deviation of each gene is equal to 1. Lastly, the expression values of each gene are scaled to [0, 1] linearly through Min-Max scaling. It is worth noting that the first step is applied to the raw datasets, while the last three steps are applied to the datasets after gene selection.

## Supporting information

**S1 Fig. Accuracy comparison between mtANN and each base classification model.** Each plot is named by query dataset. In each plot, each column represents a reference dataset. Each point is the performance of a base classification model, and points of different colors represent different gene selection methods. The red line indicates the performance of mtANN that integrates different reference datasets and gene selection methods. (A) PBMC collection. (B) Pancreas collection.

**S2 Fig. A graphical illustration of experiments.** To simulate the case where the query dataset contains unseen cell types, we alternatively remove cells belonging to one cell type shared by all the reference datasets and the query dataset, and every shared cell type will be removed once. For example, there are three reference datasets and three cell types shared between all the reference datasets and the query dataset. Thus, there will be three tests when using the three reference datasets to annotate the query dataset. In test 1, all the cells belonging to the yellow cell type in all the reference datasets are removed, so the real unseen cell type is the yellow cell type. In test 2, all the cells belonging to the blue cell type in all the reference datasets are removed, so the real unseen cell type is the blue cell type. Similarly, in test 3, all the cells belonging to the red cell type in all the reference datasets are removed, so the real unseen cell type is the red cell type. Multiple test experiments (50 tests in PBMC collection and 25 tests in Pancreas collection) are conducted by alternatively removing each shared cell type.

**S3 Fig. Performance in unseen cell-type identification.** The boxplots of the AUPRC scores of different methods in (A) PBMC collection and (B) Pancreas collection. The results with different query datasets are displayed in different panels.

**S4 Fig. Performance summary of mtANN and other compared methods in unseen cell-type identification and cell-type annotation.** The bar plot of the number of times each method ranks first in each evaluation metric. The evaluation metrics are indicated at the top of the graph and dataset collections are illustrated below the graph. Under each evaluation metric, the top 3 methods are marked with rankings.

**S5 Fig. Distribution of metric that measures cell prediction uncertainty when “10X v3” in PBMC collection is query dataset and “B cell” is the real unseen cell type.** The distribution of metric obtained from (A), ItClust (B), scGCN (enrichment) (C), scGCN (entropy) (D), scANVI are illustrated. The color of the histogram distinguishes unseen cell types from shared cell types.

**S6 Fig. Distribution of metric that measures cell prediction uncertainty when “10X v3” in PBMC collection is query dataset and “CD14+ monocyte” is the real unseen cell type.** The distribution of metric obtained from

(A), mtANN (B), scmap-clust (C), scmap-cell (D), Seurat v3 (E), ItClust (F), scGCN (enrichment) (G), scGCN (entropy) (H), scANVI are illustrated. The color of the histogram distinguishes unseen cell types from shared cell types. The black dotted line in (A) represents the subpopulations of the Gaussian mixture model fitted by mtANN. The grey solid lines in (A-D) represent the default thresholds selected by mtANN, scmap-clust, scmap-cell, and Seurat v3.

**S7 Fig. Distribution of metric that measures cell prediction uncertainty when “10X v3” in PBMC collection is query dataset and “Megakaryocyte” is the real unseen cell type**. The distribution of metric obtained from (A), mtANN

(B), scmap-clust (C), scmap-cell (D), Seurat v3 (E), ItClust (F), scGCN (enrichment) (G), scGCN (entropy) (H), scANVI are illustrated. The color of the histogram distinguishes unseen cell types from shared cell types. The black dotted line in (A) represents the subpopulations of the Gaussian mixture model fitted by mtANN. The grey solid lines in (A-D) represent the default thresholds selected by mtANN, scmap-clust, scmap-cell, and Seurat v3.

**S8 Fig. Performance in unseen cell-type identification under the default threshold.** The boxplots of the F1 scores of different methods in (A) PBMC collection and (B) Pancreas collection. The results with different query datasets are displayed in different panels.

**S9 Fig. Comparison between the true proportion of unseen cell types and the proportion of unassigned cells predicted by each method**. Dot plots of all tests (75 tests) when all datasets in the PBMC collection and Pancreas collection are query datasets respectively. In each dot plot, the *x*-axis is the true proportion and the *y*-axis is the proportion of unassigned cells predicted by each method, and the different colors of the dots represent the different methods. The black solid line is the line of *y* = *x*. The Pearson correlations between true proportion and proportion of unassigned cells predicted by each method are reported.

**S10 Fig. Cell-type annotation performance with the real proportion of unseen cell types as a threshold**. The boxplots of the accuracy of different methods in (A) PBMC collection and (B) Pancreas collection. The results with different query datasets are displayed in different panels.

**S11 Fig. Similarity of the annotation labels of each method to the real label.** The heatmap of Pearson correlations between cell-type proportions of the true cell-type label and annotation labels predicted by each method in (A) PBMC collection and (B) Pancreas collection. Columns of the heatmap represent the 50 tests in PBMC collection and the 25 tests in Pancreas collection.

**S12 Fig. Cell-type annotation performance with the default threshold.** The boxplots of the accuracy of different methods in (A) PBMC collection and (B) Pancreas collection. The results with different query datasets are displayed in different panels.

**S13 Fig. Performance comparison between mtANN using the real proportion of unseen cell types and mtANN using the default threshold**. The boxplots of the accuracy of mtANN using the real proportion of unseen cell types and mtANN using the default threshold in (A) PBMC collection and (B) Pancreas collection. The results with different query datasets are displayed in different panels.

**S1 Text. Supplementary notes of mtANN.** There are algorithm, details in the Modules I-IV of mtANN, methods for benchmark and performance assessment.

**S1 Table. Terms and notations.**

**S2 Table. Gene selection threshold settings.**

**S3 Table. The cell types and cell numbers of each dataset in PBMC collection.**

**S4 Table. The cell types and cell numbers of each dataset in Pancreas collection.**

**S5 Table. The query datasets, references and the real unseen cell type of each experiment test in PBMC collection.**

**S6 Table. The query datasets, references and the real unseen cell type of each experiment test in Pancreas collection.**

## Acknowledgments

We acknowledge all members of the Xiao-Fei Zhang laboratory for helpful suggestions.

## Author Contributions

Conceptualization: Yi-Xuan Xiong, Luonan Chen, Xiao-Fei Zhang.

Data curation: Yi-Xuan Xiong, Meng-Guo Wang.

Formal analysis: Yi-Xuan Xiong.

Funding acquisition: Luonan Chen, Xiao-Fei Zhang. Investigation: Meng-Guo Wang, Xiao-Fei Zhang.

Methodology: Yi-Xuan Xiong, Xiao-Fei Zhang. Software: Yi-Xuan Xiong, Meng-Guo Wang.

Supervision: Luonan Chen, Xiao-Fei Zhang. Visualization: Yi-Xuan Xiong.

Writing - original draft: Yi-Xuan Xiong.

Writing - review & editing: Yi-Xuan Xiong, Luonan Chen, Xiao-Fei Zhang.

