## Supplementary material for "Cell-type Annotation with Accurate Unseen Cell-type Identification Using Multiple References": S1_Text.pdf

### Supplementary Information for the article: "Cell-type Annotation with Accurate Unseen Cell-type Identification Using Multiple References"

Yi-Xuan Xiong[1,2], Meng-Guo Wang[1,2], Luonan Chen[3,4,5,6,*], Xiao-Fei Zhang[1,2,*]

[1] *School of Mathematics and Statistics, Central China Normal University, Wuhan 430079, China*

[2] *Hubei Key Laboratory of Mathematical Sciences, Central China Normal University, Wuhan 430079, China*

[3] *State Key Laboratory of Cell Biology, Shanghai Institute of Biochemistry and Cell Biology, Center for Excellence in Molecular Cell Science, Chinese Academy of Sciences, Shanghai 200031, China*

[4] *School of Life Science and Technology, ShanghaiTech University, Shanghai 201210, China*

[5] *Key Laboratory of Systems Health Science of Zhejiang Province, Hangzhou Institute for Advanced Study, University of Chinese Academy of Sciences, Chinese Academy of Sciences, Hangzhou 310024, China*

[6] *Guangdong Institute of Intelligence Science and Technology, Hengqin, Zhuhai, Guangdong 519031, China*

 (Zhang XF), (Chen L)

#### The workflow of mtANN

**Algorithm 1** The entire workflow of mtANN

**Input:**

M well-labeled reference datasets: $\{(X^{r_i}, Y^{r_i})\}_{i=1}^{M}$

The query dataset: $X^q$

Count vector: $L$

**for** $i = 1$ to $M$ **do**:

**for** j= 1 to 8 **do**:

**Module I:** Gene selection;

select gene set $G^{r_{ij}}$ on $X^{r_i}$ with $j$-th gene selection method

$X^{r_{ij}} = X^{r_i}[:, G^{r_{ij}} \cap G^q]$

$X^{q_{ij}} = X^q[:, G^{r_{ij}} \cap G^q]$

construct dataset pair $(X^{r_{ij}}, Y^{r_i}, X^{q_{ij}})$ with preprocessed $X^{r_{ij}}$ and $X^{q_{ij}}$

**Module II:** Deep classification model training;

$\hat{X}^{r_{ij}} = D^{ij}\left(E^{ij}(X^{r_{ij}})\right)$

$\hat{X}^{q_{ij}} = D^{ij}\left(E^{ij}(X^{q_{ij}})\right)$

$\hat{P}^{r_{ij}} = softmax\left(C^{ij}\left(E^{ij}(X^{r_{ij}})\right)\right)$

$\mathcal{L}_{re} = \frac{1}{n^{r_i}p^{ij}} \| \hat{X}^{r_{ij}} - X^{r_{ij}} \|_F^2 + \frac{1}{n^q p^{ij}} \| \hat{X}^{q_{ij}} - X^{q_{ij}} \|_F^2$

$\mathcal{L}_{ce} = -\frac{1}{n^{r_i}} \sum_{c=1}^{n^{r_i}} \sum_{k \in K} \mathbf{1}_{\left[Y_c^{r_{ij}} = k\right]} \log \hat{P}_{ck}^{r_{ij}}$

initialize encoder $E^{ij}$ and decoder $D^{ij}$ with loss function $\mathcal{L}_{re}$

finetune $E^{ij}$, $D^{ij}$ and train classifier $C^{ij}$ simultaneously with loss function $\mathcal{L}_{ce} + \lambda \mathcal{L}_{re}$

**end for**

**end for**

**Module III:** Query dataset annotation;

with each base classification model $\left\{\{E^{ij}, C^{ij}\}_{i=1}^{M}\right\}_{j=1}^{8}$,

$\hat{P}^{q_{ij}} = softmax\left(C^{ij}\left(E^{ij}(X^{q_{ij}})\right)\right)$

for cell $c$, $\hat{Y}_c^{q_{ij}} = \underset{k \in K}{\operatorname{argmax}} \hat{P}_{ck}^{q_{ij}}$

obtain consensus annotation $\hat{Y}^q$ based on majority voting with $\hat{Y}_c^q = \underset{k \in K}{\operatorname{argmax}} \frac{\sum_{i=1}^{M} \sum_{j=1}^{8} \mathbf{1}_{\left[\hat{Y}_c^{q_{ij}} = k\right]}}{L_k}$

**Module IV:** Metrics for unseen cell identification;

for cell $c$:

calculate intra-model measurement $\mathrm{m}_c^{(1)} = \frac{1}{8\mathrm{M}} \sum_{i,j} H\left(\hat{P}_c^{q_{ij}}\right)$

$Q_{ck}^{(2)} = \frac{\sum_{i,j} \hat{P}_{ck}^{q_{ij}}}{L_k}$

$Q_{ck}^{(3)} = \frac{\sum_{i,j} \mathbf{1}_{\left[\hat{Y}_c^{q_{ij}}=k\right]}}{L_k}$

$\tilde{Q}^{(2)} = Q_c^{(2)}/sum\left(Q_c^{(2)}\right)$

$\tilde{Q}^{(3)} = Q_c^{(3)}/sum\left(Q_c^{(3)}\right)$

calculate inter-model measurement $m_c^{(2)} = H\left(\tilde{Q}_c^{(2)}\right)$

calculate inter-model measurement $m_c^{(3)} = H\left(\tilde{Q}_c^{(3)}\right)$

**for** l= 1 to 3 **do**:

$\bar{m}^{(l)} = \left(m^{(l)} - \min\left(m^{(l)}\right)\right)/\left(\max\left(m^{(l)}\right) - \min\left(m^{(l)}\right)\right)$

**end for**

obtain metric for unseen cell-type identification $m_c = \frac{\bar{m}_c^{(1)}+\bar{m}_c^{(2)}+\bar{m}_c^{(3)}}{3}$

**Module V:** Data-driven method for default threshold selection;

**for** S= 1 to 5 **do**:

construct Gaussian mixture model: $p(m) = \sum_{s=1}^{S} \pi_s N(\mu_s, \sigma_s^2)$

calculate the AIC of the model, denoted by $\mathrm{AIC_S}$

**end for**

$S^* = \arg\min_S AIC_S$

**if** $S^*$ == 1:

no unassigned cells

**else**:

obtain cell groups $\mathrm{g^q}$ according to the posterior probability

calculate the mean of uncertainty metric within each group $\{\tilde{m}_1, \cdots, \tilde{m}_{S^*}\}$

**end if**

**if** $\max_{s\in\{1,\dots,S^*\}} \tilde{m}_S < 0.6$:

unassigned cells set: cells where $g^q == arg \max_s \tilde{m}_S$

**else:**

unassigned cells set: cells where $g^q == s$ if $\tilde{m}_S \geq 0.6$

**end if**

**Output:**

$\hat{Y}^q$: the annotated cell type labels of cells in the query dataset

$U$: the set of unassigned cells

### Details in the workflow of mtANN

#### Module I： Gene selection

Eight gene selection methods are adopted by mtANN to select genes, including Limma, Bartlett's test, Kolmogorov-Smirnov test, Chi-squared test, Bimodality index, Gini index, Dispersion and Variance-stabilizing transformation. The first five gene selection methods (i.e., Limma, Bartlett's test, Kolmogorov-Smirnov test, Chi-squared test and Bimodality index) first select genes for each cell type separately by comparing the cells belonging to that cell type against the rest cells from all other cell types, and then integrate all these genes, while the last three gene selection methods (i.e., Gini index, Dispersion and Variance-stabilizing transformation) select genes based on the entire dataset. It is worth noting that we select genes on the raw data after library size normalization and preprocess the data after gene selection. The following is a brief introduction to eight gene selection methods. For specific gene selection threshold settings, please refer to Table S2.

- Limma. This method identifies genes based on linear models and empirical Bayes methods. For each cell type, differential expression analysis is implemented by using the functions *lmFit*, *eBayes* and *topTable* in sequence in the *limma* package[1]. The significant genes are extracted as differentially expressed genes for one cell type.
- Bartlett's test. This method selects genes based on variance. For each cell type, the null hypothesis for each gene is that the variance of the cells belonging to this cell type is the same as the variance of the rest. This procedure is implemented by the *bartlett.test* function in R. The significant genes are extracted as differential variable genes for one cell type.
- Kolmogorov-Smirnov test. This method selects genes based on distribution. For each cell type, the null hypothesis for each gene is that the cumulative distribution function of the rest cells is not greater than the cumulative distribution function of the cells from the selected cell type. This procedure is implemented by the *ks.test*

function in R. The significant genes are extracted as differentially distributed genes for one cell type.

- Chi-squared test. This method selects genes based on the proportion of expressed cells. Here, a gene is considered to be expressed in a cell if its expression level is greater than 1. For each cell type, the null hypothesis for each gene is that the gene expressed in the cells from this cell type is not dependent on this cell type. This procedure is implemented by the *chisq.test* function in R. The significant genes are extracted as differentially proportioned genes for one cell type.
- Bimodality index. This method identifies genes with bimodal expression patterns. For each cell type, the bimodality index (BI)[2, 3] of each gene can be calculated by

$$BI = \sqrt{\pi(1-\pi)}\frac{|\mu_1 - \mu_2|}{\sqrt{\pi\sigma_1^2 + (1-\pi)\sigma_2^2}} \tag{1}$$

  where $\mu_1$ and $\sigma_1$ are the mean and the standard deviation of the expression of the cells from the cell type, $\mu_2$ and $\sigma_2$ are the mean and the standard deviation of the expression of the rest cells, and $\pi$ is the proportion of the cells from the cell type. The genes with the largest BI values are extracted as high BI genes for one cell type.
- Gini index. This method has advantages in identifying rare cell-type-specific genes[4]. For each gene, the Gini index is defined as twice the area between the Lorenz curve and the diagonal. This procedure is implemented by the *calGini* function of the *giniclust3* package in Python, which also includes a normalizing procedure to remove bias for low-expressed genes[5]. The genes with the largest Gini index values are extracted as high Gini genes.
- Dispersion. This method selects genes based on dispersion. For each gene, dispersion is defined as the ratio between the variance and the mean of expression across all the cells. The genes with the largest dispersion values are extracted as highly variable genes. This procedure is implemented by the *FindVariableFeatures* function of the *Seurat* package in R with the argument *selection.method="disp"*.[6]

- Variance-stabilizing transformation. This method selects genes based on variance after correction by variance-stabilizing transformation (vst). The mean-variance relationship is fitted by a LOESS regression model, and then the expression value of each gene is standardized by the observed mean and expected variance. The vst-corrected variance for each gene is calculated based on the standardized expression values. The genes with the largest vst-corrected variances are extracted as highly variable genes. This procedure is implemented by the *FindVariableFeatures* function of the *Seurat* package in R with the argument *selection.method=“vst”*.[6]

**Module II： Deep classification model training**

- Neural network architecture

  The embedding component is composed of two fully connected layers. The first layer consists of 128 neurons, and the second layer consists of 32 neurons. Batch normalization and the rectified linear unit (ReLU) function are applied to the first layer. The number of neurons in the input layer is determined by the number of genes of the corresponding dataset pair. The number of neurons in the linear classifier layer is determined by the number of cell types observed in the reference subset of the corresponding dataset pair.

- Hyperparameter settings

  The parameters of the embedding component, the decoder component and the linear classifier layer are updated by Adam with the learning rate of 0.01. The batch size of $X^{r_{ij}}$ is set as $\left[\left(\max_{k\in K}\sum_{c=1}^{n^{r_i}} 1_{\left[Y_c^{r_i}=k\right]}\right)/10\right]$ and the batch size of $X^{q_{ij}}$ is set as $\min([n^q/10], 600)$ for dataset pair $(X^{r_{ij}}, Y^{r_i}, X^{q_{ij}})$, where $[\cdot]$ indicates rounding up to an integer. The training process is implemented with the epoch of 30.

- Initialization

  Before training the deep classification model, we initialize the parameters of the embedding component and the decoder component by training an autoencoder. For dataset pair $(X^{r_{ij}}, Y^{r_i}, X^{q_{ij}})$, we take the gene expression of cells both from the reference subset and the query subset as input and train the autoencoder to reconstruct the gene expression well. The reconstruction loss can be formulated as

$$\mathcal{L}_{re} = \frac{1}{n^{r_i} p^{ij}} \|\hat{X}^{r_{ij}} - X^{r_{ij}}\|_F^2 + \frac{1}{n^q p^{ij}} \|\hat{X}^{q_{ij}} - X^{q_{ij}}\|_F^2,$$

and the optimization problem for initialization can be written as

$$\min_{E^{ij}, D^{ij}} \quad \mathcal{L}_{re}. \tag{2}$$

The parameters of the resulting embedding component and decoder component are then used as the initial parameters for training the classification model.

**Module III : Query dataset annotation**

With each base classification model $\left\{\{E^{ij}, C^{ij}\}_{i=1}^{M}\right\}_{j=1}^{8}$, we calculate prediction probability with $\hat{P}^{q_{ij}} = softmax\left(C^{ij}\left(E^{ij}(X^{q_{ij}})\right)\right)$. Thus, we can obtain base predictions $\left\{\{\hat{Y}^{q_{ij}}\}_{i=1}^{M}\right\}_{j=1}^{8}$ and obtain a consensus annotation $\hat{Y}^q$ based on majority voting. However, if a cell is predicted to be of different types by the same number of base classification models, we determine the final label according to the maximum sum of their corresponding prediction probabilities. For example, cell $c$ is annotated $4M$ times as cell type $k$ and $\tilde{k}$ seperately. Calculate

$$\Sigma(\hat{P}_{ck}^{q_{ij}}) \tag{3}$$

where $\hat{Y}_c^{q_{ij}} = k$, and calculate

$$\Sigma(\hat{P}_{c\tilde{k}}^{q_{\tilde{i}\tilde{j}}}) \tag{4}$$

where $\hat{Y}_c^{q_{\tilde{i}\tilde{j}}} = \tilde{k}$. Thus, the final annotation of cell $c$ is $k$ if equation (3) is greater than equation (4) or vice versa.

**Module IV : Metrics for unseen cell identification**

With all the base classification models, we can obtain a series of prediction probabilities $\left\{\{\hat{P}^{q_{ij}}\}_{i=1}^{M}\right\}_{j=1}^{8}$. For cell $c$, we integrate all the prediction probabilities into a matrix

| | Cell type 1 | Cell type 2 | ⋯ | Cell type $K$ |
|---|---|---|---|---|
| Model 1 | $\hat{P}_{c1}^{q_{11}}$ | $\hat{P}_{c2}^{q_{11}}$ | ⋯ | $\hat{P}_{cK}^{q_{11}}$ |
| Model 2 | $\hat{P}_{c1}^{q_{12}}$ | $\hat{P}_{c2}^{q_{12}}$ | ⋯ | $\hat{P}_{cK}^{q_{12}}$ |
| ⋮ | ⋮ | ⋮ | ⋮ | ⋮ |
| Model $8M$ | $\hat{P}_{c1}^{M8}$ | $\hat{P}_{c2}^{q_{M8}}$ | ⋯ | $\hat{P}_{cK}^{q_{M8}}$ |

Then, we calculate entropy of each row which can be denoted as $H(\hat{P}^{q_{ij}}), i = 1 \cdots M, j = 1 \cdots 8$. Finally, the intra-model measurement $m^{(1)} = \frac{1}{8M}\sum_{i,j} H\left(\hat{P}^{q_{ij}}\right)$.

For the inter-model measurement $m^{(2)}$, we sum up the prediction probabilities by column and divide it by the number of reference subsets that containing the corresponding cell type which is denoted as $Q_c^{(2)}$:

| Cell type 1 | Cell type 2 | ⋯ | Cell type $K$ |
|---|---|---|---|
| $\frac{\left(\hat{P}_{c1}^{q_{11}} + \hat{P}_{c1}^{q_{12}} + \cdots + \hat{P}_{c1}^{M8}\right)}{L_1}$ | $\frac{\left(\hat{P}_{c2}^{q_{11}} + \hat{P}_{c2}^{q_{12}} + \cdots + \hat{P}_{c2}^{M8}\right)}{L_2}$ | ⋯ | $\frac{\left(\hat{P}_{cK}^{q_{11}} + \hat{P}_{cK}^{q_{12}} + \cdots + \hat{P}_{cK}^{M8}\right)}{L_K}$ |

For all the cells, $Q^{(2)}$ is transformed into a probability matrix $\widetilde{Q}^{(2)}$ by dividing each value by the row sum. Finally, the inter-model measurement $m_c^{(2)} = H\left(\widetilde{Q}_c^{(2)}\right)$.

For the inter-model measurement $m^{(3)}$, we obtain all the base prediction label of cell $c$ according to prediction probabilities $\left\{\left\{\hat{P}^{q_{ij}}\right\}_{i=1}^{M}\right\}_{j=1}^{8}$

| | |
|---|---|
| Model 1 | $\hat{Y}_c^{q_{11}}$ |
| Model 2 | $\hat{Y}_c^{q_{12}}$ |
| ⋮ | ⋮ |
| Model $8M$ | $\hat{Y}_c^{q_{M8}}$ |

Then, $Q_{ck}^{(3)} = \frac{\sum_{i,j} \mathbf{1}_{\left[\hat{Y}_c^{q_{ij}}=k\right]}}{L_k}$ represents the integrated result for this measure. We also transform $Q^{(3)}$ into a probability matrix $\widetilde{Q}^{(3)}$ by dividing each value by the row sum. Finally, the inter-model measurement $m_c^{(3)} = H\left(\widetilde{Q}_c^{(3)}\right)$.

#### Methods for benchmark

We compare mtANN with seven cell-type annotation methods including scmap-clust[7], scmap-cell[7], Seurat v3[6], ItClust[8], scGCN (enrichment)[9], scGCN (entropy)[9], and scANVI[10]. scmap-clust, scmap-cell, and Seurat v3 can be applied to cell-type annotation based on multiple reference datasets. For the other four methods which are designed to annotate cell types based on a single reference dataset, we directly combine multiple well-annotated datasets as reference datasets in order to use consistent datasets with other methods. The following is a brief introduction to the steps of these methods to identify unseen cells.

- Seurat v3. We first integrate multiple reference datasets using the function *IntegrateData* in Seurat under the default parameters, and then perform cell-type annotation with function *TransferData*. In addition, Seurat v3 identifies "unassigned" cells according to the predicted probability. The smaller the probability, the more likely the corresponding cell is of an unseen cell type. The default threshold for unseen cell type identification provided by Seurat v3 is the 20-$th$ percentile of the probabilities.
- scmap. It projects query cells onto cell types or individual cells, and these two types of methods are denoted as scmap-clust and scmap-cell, respectively. We perform cell-type annotation based on multi-reference datasets according to the vignette provided by scmap. In addition, scmap-clust and scmap-cell identify "unassigned" cells according to the integrative predicted probability. The smaller the probability, the more likely the corresponding cell is unseen. For identifying unseen cell types, the default thresholds provided by scmap-clust and scmap-cell are 0.7 and 0.5.
- ItClust. It identifies unseen cell types using a confidence score defined by calculating the similarity between clusters and cell types in the reference dataset. The smaller the similarity, the more likely the corresponding cluster is of an unseen cell type.
- scGCN. It proposes two metrics to identify unseen cell types, which are entropy and enrichment. Identifying unseen cell types with different metrics is regarded as two methods: scGCN (enrichment) and scGCN (entropy). The smaller the enrichment, the more likely the corresponding cell is of an unseen

cell type. Conversely, the greater the entropy, the more likely the corresponding cell is of an unseen cell type.

- scANVI. The unseen cells are identified based on the prediction probability. The smaller the probability, the more likely the corresponding cell is of an unseen cell type.

For all the methods, the data preprocessing procedures followed their original manuscript and all the parameters are the default values.

#### Performance assessment

We use the area under the precision-recall curve (AUPRC) score to evaluate the performance of unseen cell type identification metrics of different methods. The x-axis and y-axis of the precision-recall curve are $recall$ and $precision$ which can be calculated as below:

$$recall = \frac{TP}{TP + FN}, \tag{5}$$

$$precision = \frac{TP}{TP + FP}, \tag{6}$$

where $TP$ is the number of cells identified as “unassigned” that belong to the truly unseen cell type. $FP$ represents the number of cells identified as “unassigned” that are not belong to the truly unseen cell type, and $FN$ means the number of cells of the truly unseen cell type that are not identified as “unassigned”.

At a fixed threshold, we use the F1 score to compare the accuracy of each method in identifying unseen cell types in the query dataset. F1 score can be calculated as below:

$$F1\ score = 2 \cdot \frac{precision \cdot recall}{precision + recall}. \tag{7}$$

To compare the performance of mtANN with other methods in annotating the entire query dataset, we use accuracy which is defined as the proportion of correct predictions in the query dataset. Accuracy can be calculated as below:

$$Accuracy = \frac{1}{n}\sum_{i=1}^{n} 1_{[\hat{y}_i = y_i]}, \tag{8}$$

where $n$ is the number of query cells, $y_i$ is the true cell-type of $i$-th cell where the cell belongs to the real unseen cell type is labeled as "unassigned", and $\hat{y}_i$ is the predicted cell type of $i$-th cell where the cell labeled as "unassigned" belongs to the predicted unseen cell type.
